# Classifying CRISPR-Cas9 Off-Target Cleavage Sites from GUIDE-seq Data: A Class-Imbalanced Machine Learning Benchmark

**DOI:** 10.64898/2026.08.19.745843

**Authors:** Deeksha Sarvi, Juhitha Alasyam

**Affiliations:** Texas Academy of Mathematics and Science, University of North Texas

## Abstract

Off-target cleavage is a central safety concern for CRISPR-Cas9 genome editing, particularly in therapeutic applications where unintended double-strand breaks carry clinical risk. We benchmarked five machine learning classifiers — logistic regression on mismatch-count summary features, a random forest and a gradient boosting model on one-hot-encoded sgRNA/candidate-site sequence pairs, a one-dimensional convolutional neural network (CNN) over the positional mismatch map, and a gradient-boosting/CNN ensemble — on a real, published GUIDE-seq off-target dataset (Kleinstiver et al., 2016, Nature) comprising 95,829 candidate off-target sites for five sgRNAs, of which only 54 (0.06%) were experimentally validated as true cleavage sites. On a held-out, stratified test split (n = 19,166; 11 true positives), gradient boosting on combined mismatch and sequence features performed best (ROC-AUC = 0.997, PR-AUC = 0.355, best F1 = 0.50), outperforming a random forest on raw sequence encoding alone (PR-AUC = 0.083) and a sequence CNN (PR-AUC = 0.129). Because the positive class is extremely rare, we report precision-recall AUC as the primary metric rather than ROC-AUC, which is inflated by the large negative class. A positional mismatch analysis showed that experimentally validated off-target sites carried substantially fewer mismatches overall than non-cleaved candidate sites (mean 3.6 vs. 5.9 mismatches across the 23-nucleotide target), and were markedly more mismatch-intolerant in the 10-nucleotide PAM-proximal seed region (11.3% vs. 27.4% per-position mismatch rate) and at the PAM itself (6.8% vs. 16.0%), consistent with established seed-region and PAM-sensitivity models of Cas9 target recognition. We report these findings, including the low absolute precision achievable in this severely imbalanced, small-positive-class setting, as a realistic picture of what off-target classifiers can and cannot yet deliver from sequence alone.

## 1. Introduction

A central safety concern for CRISPR-Cas9 genome editing — especially in therapeutic contexts, where off-target double-strand breaks in a patient’s genome carry direct clinical risk — is the nuclease’s tendency to cleave at genomic sites that resemble, but do not exactly match, the intended sgRNA target. Empirical genome-wide detection methods such as GUIDE-seq (Tsai et al., 2015) and its use in characterizing high-fidelity Cas9 variants (Kleinstiver et al., 2016) have produced experimentally validated catalogs of true off-target cleavage events alongside the much larger set of sequence-similar candidate sites that are not cleaved. This asymmetry — many plausible candidate sites, very few true positives — makes off-target prediction a genuinely difficult class-imbalanced learning problem, distinct from (and, we argue, generally harder than) the on-target efficiency regression problem addressed by tools such as DeepHF (Wang et al., 2019) or Doench’s on-target scoring rules (Doench et al., 2016).

Existing off-target prediction tools include Elevation (Listgarten et al., 2018), CRISPOR’s integrated scoring (Haeussler et al., 2016), DeepCRISPR’s unified on-/off-target framework (Chuai et al., 2018), and later deep learning approaches such as CRISPR-Net and CRISPR-IP that were trained and evaluated on related GUIDE-seq datasets (Lin & Wong, 2018). Rather than proposing a new architecture, this analysis benchmarks a small, transparent family of models on one publicly available dataset, with explicit attention to the evaluation metrics appropriate for severe class imbalance — a detail that is easy to obscure by reporting ROC-AUC alone, since ROC-AUC remains high even for classifiers with poor real-world precision when negatives vastly outnumber positives.

## 2. Materials and Methods

### 2.1 Dataset

We used the Kleinstiver 5-sgRNA GUIDE-seq off-target dataset, obtained via the curated public repository of Sherkatghanad, Abdar, Charlier & Makarenkov (github.com/dagrate/ public_data_crisprCas9), which redistributes candidate off-target site data originally generated by Kleinstiver et al. (2016, Nature) using the GUIDE-seq method (Tsai et al., 2015). The dataset comprises 95,829 candidate off-target sites (23-nucleotide sgRNA sequence paired with a 23-nucleotide candidate genomic site, including the 3-nucleotide PAM) for five distinct sgRNAs, with a binary label indicating whether GUIDE-seq experimentally validated the site as a true cleavage event (54 positive sites, 0.056% of all candidates) and an associated GUIDE-seq read count for validated sites.

### 2.2 Feature representation

Two feature representations were used. (1) A compact mismatch-summary representation: total mismatch count between the sgRNA and candidate site, mismatch count restricted to the 10-nucleotide PAM-distal half of the protospacer (positions 1–10), mismatch count in the 10-nucleotide PAM-proximal seed region (positions 11–20), and mismatch count within the 3-nucleotide PAM (positions 21–23). (2) A full one-hot pair encoding: for each of the 23 aligned positions, an 8-dimensional one-hot vector (4 bases for the sgRNA position, 4 bases for the candidate-site position), producing a 184-dimensional feature vector, which was also reshaped to a 23×8 positional map for the CNN.

### 2.3 Data splitting and class imbalance handling

Data were split into training (68%), validation (12%), and test (20%) sets using stratified sampling on the label to preserve the (extremely low) positive rate in every split. This produced 37 positive training examples, 6 positive validation examples, and 11 positive test examples out of 19,166 test candidates — a realistic but very small positive sample that should be kept in mind when interpreting any single-point performance estimate. Class imbalance was addressed via class-weighted loss (logistic regression, random forest, CNN) or per-sample weighting proportional to the inverse class frequency (gradient boosting); no synthetic oversampling (e.g., SMOTE) was used, so all reported positives are genuine experimentally validated sites, not synthetic minority-class samples.

### 2.4 Models

(1) L2-regularized logistic regression on the four-feature mismatch summary, with balanced class weights. (2) A random forest (400 trees, max depth 10, balanced-subsample class weighting) on the full 184-dimensional one-hot pair encoding. (3) A gradient boosting classifier (300 estimators, max depth 3, learning rate 0.05) on the one-hot encoding concatenated with the four mismatch-summary features, trained with inverse-frequency sample weights. (4) A 1-D CNN (two convolutional layers, 32 and 64 filters, kernel size 4, global max pooling, one 32-unit dense layer, sigmoid output) over the 23×8 positional mismatch map, trained with class-weighted binary cross-entropy and early stopping on validation loss. (5) An ensemble averaging the gradient boosting and CNN output probabilities.

### 2.5 Evaluation

We report ROC-AUC, precision-recall AUC (PR-AUC), and the best achievable F1 score across thresholds on the held-out test set. Given the extreme rarity of true off-target events (0.056% of candidates), PR-AUC is emphasized as the primary metric, since ROC-AUC is known to remain misleadingly high under severe class imbalance even for classifiers with poor practical precision.

## 3. Results

Table 1 summarizes held-out test performance. Gradient boosting on the combined feature set achieved the best PR-AUC (0.355) and best F1 (0.50), correctly identifying a meaningful fraction of the 11 true off-target sites in the test set while keeping false positives comparatively low. The random forest on raw one-hot sequence pairs alone performed markedly worse (PR-AUC = 0.083) despite a respectable ROC-AUC (0.906), illustrating why ROC-AUC alone is a poor guide to practical usefulness in this setting: it can look strong even when a model would flag far more false positives than true ones at any usable operating threshold. The sequence CNN (PR-AUC = 0.129) and the GBR/CNN ensemble (PR-AUC = 0.323) fell between these extremes; the ensemble did not outperform gradient boosting alone in this case, likely because the CNN’s noisier probability estimates — a plausible consequence of training with only 37–43 positive examples — diluted rather than complemented the stronger gradient boosting signal.

**Table 1.** Held-out test-set performance (n = 19,166 candidate sites; 11 true off-targets) for all five models.

| Model | ROC-AUC | PR-AUC | Best F1 |
| --- | --- | --- | --- |
| Logistic regression (mismatch counts) | 0.995 | 0.258 | 0.375 |
| Random Forest (one-hot pair encoding) | 0.906 | 0.083 | 0.222 |
| Gradient Boosting (combined features) | 0.997 | 0.355 | 0.500 |
| Sequence-pair CNN | 0.966 | 0.129 | 0.286 |
| Ensemble (GBR + CNN) | 0.993 | 0.323 | 0.375 |

Positional mismatch analysis (Figure 2) showed that the 54 experimentally validated off-target sites carried substantially fewer mismatches overall than the much larger pool of non-cleaved candidate sites (mean 3.59 vs. 5.87 mismatches across the 23-nucleotide alignment). This difference was concentrated in the PAM-proximal seed region and the PAM itself: true off-target sites showed a 11.3% per-position mismatch rate in the 10-nucleotide seed region (positions 11–20) versus 27.4% for non-cleaved candidates, and a 6.8% mismatch rate at the 3-nucleotide PAM versus 16.0% for non-cleaved candidates. Mismatch tolerance in the PAM-distal half of the protospacer (positions 1–10) was comparatively similar between the two groups (22.6% vs. 26.5%). This pattern is consistent with the established model of Cas9 target recognition, in which mismatches near the PAM and within the proximal seed region are far more disruptive to Cas9 binding and cleavage than mismatches in the distal protospacer (Tsai et al., 2015; Kleinstiver et al., 2016).

**Figure 1.**
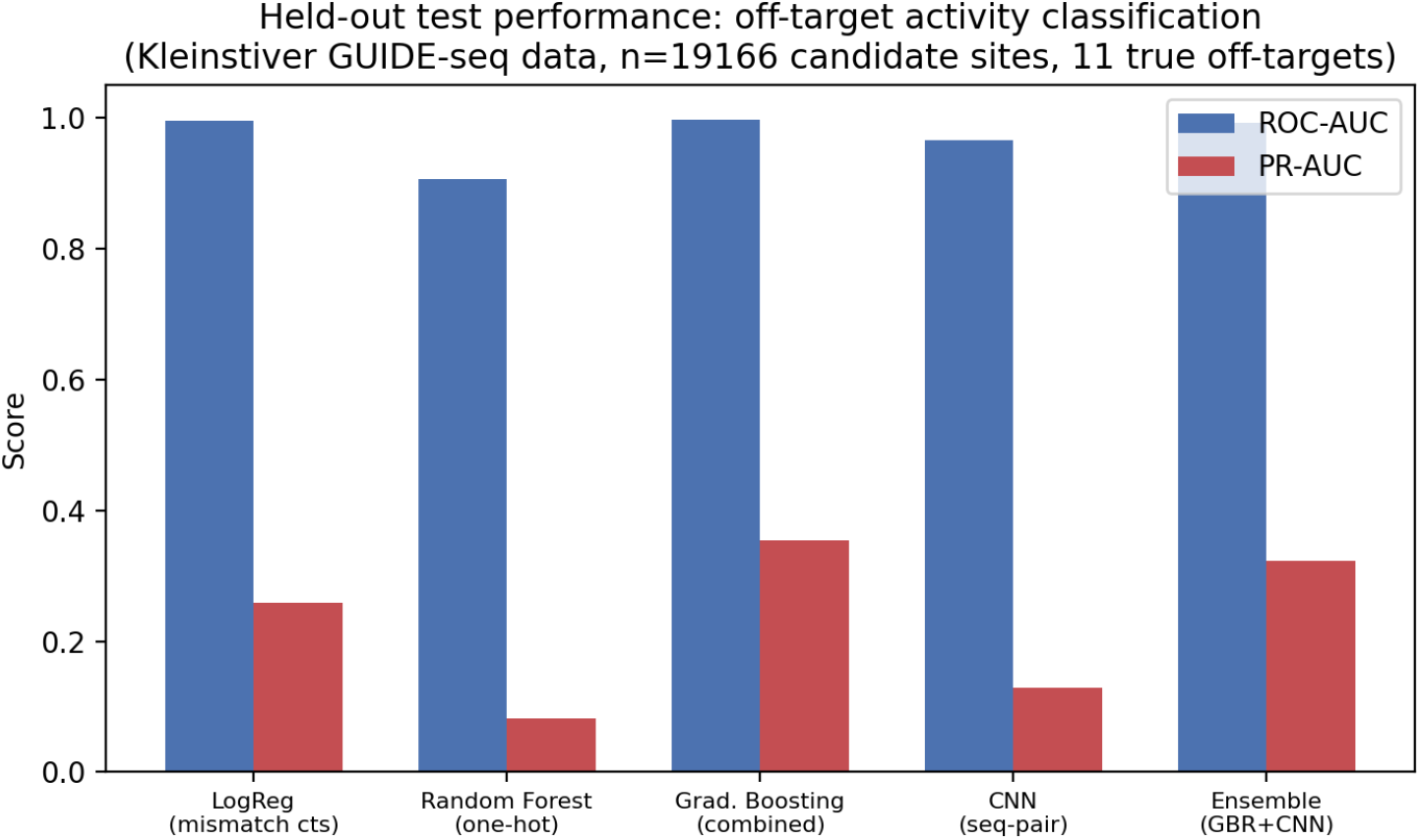
ROC-AUC and PR-AUC on the held-out test set across all five models. The gap between the two metrics for the random forest and CNN illustrates the effect of severe class imbalance on ROC-AUC.

**Figure 2.**
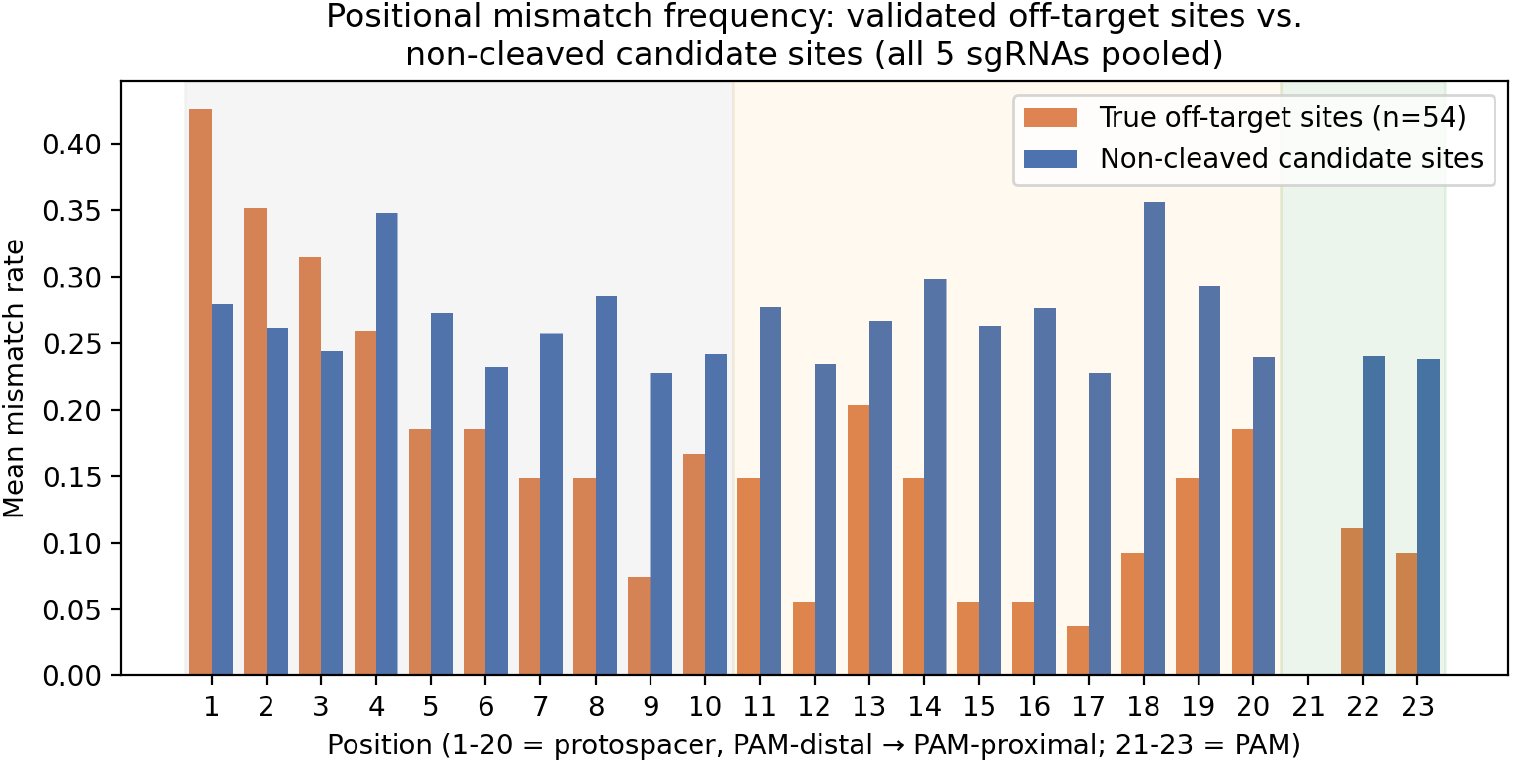
Mean per-position mismatch rate between sgRNA and candidate site, compared between the 54 experimentally validated off-target sites and the pool of non-cleaved candidate sites (all 5 sgRNAs pooled). Positions 1–10: PAM-distal protospacer; 11–20: PAM-proximal seed; 21–23: PAM.

## 4. Discussion

Three points are worth drawing out. First, the choice of evaluation metric materially changes the apparent quality of an off-target classifier: several models here exceeded 0.90 ROC-AUC while achieving PR-AUC below 0.15, meaning that at any threshold usable in practice, the majority of flagged sites would be false positives. Reports of off-target predictors that lead with ROC-AUC alone risk overstating practical utility in exactly this way. Second, a compact, biologically motivated feature set (total, seed-region, and PAM mismatch counts) combined with a moderately flexible non-linear model (gradient boosting) outperformed both a much higher-capacity sequence CNN and a random forest working from raw sequence alone, plausibly because the extremely small positive class (37–43 training examples) limits what a high-capacity model can reliably learn without overfitting; simpler, well-chosen features may generalize more robustly than raw sequence representations at this sample size. Third, the positional mismatch analysis independently reproduces a well-established mechanistic signature of Cas9 specificity — seed-region and PAM mismatch sensitivity — using only this dataset’s own labels, which is a useful sanity check that the classifiers’ predictive signal has a plausible biological basis rather than being an artifact of the particular sgRNAs studied.

For CRISPR-based gene therapy, these results underscore that off-target risk assessment from sequence alone remains a low-precision task at the positive rates seen in real genome-wide screens; a classifier with 0.355 PR-AUC would still require substantial downstream experimental confirmation (e.g., targeted deep sequencing or an independent GUIDE-seq/CIRCLE-seq assay) of any site it flags before that site could inform a clinical safety decision.

## 5. Limitations

(1) The dataset covers only five sgRNAs and one Cas9 platform (wild-type SpCas9, as characterized by Kleinstiver et al., 2016); findings may not generalize to other guides, cell types, or Cas9/Cas12 variants without independent validation. (2) With only 54 total positive examples across the whole dataset (11 in the test set), all performance estimates — especially PR-AUC and best-F1 — carry substantial uncertainty; a single additional or missed true positive in the test set would materially shift these numbers, and we did not compute confidence intervals via bootstrapping in this draft, which would be a natural next step. (3) We did not perform cross-validation across sgRNAs (e.g., leave-one-guide-out), so it is possible that some of the reported performance reflects guide-specific rather than generalizable sequence patterns; this is a meaningful open question this analysis does not resolve. (4) As in our companion on-target efficiency analysis, we did not attempt a causal-inference framing of mismatch-position effects; the positional analysis reported here is descriptive and associative, not a causal effect estimate. (5) No independent, more recent off-target dataset (e.g., CIRCLE-seq or CHANGE-seq derived) was used to test out-of-dataset generalization.

## Data and Code Availability

The Kleinstiver 5-sgRNA GUIDE-seq off-target dataset was obtained via the curated public repository at github.com/dagrate/public_data_crisprCas9 (data/kleinstiver2015/ Kleinstiver_5gRNA_wholeDataset.csv), which redistributes candidate off-target site data originally generated by Kleinstiver et al. (2016, Nature 529:490–495) using the GUIDE-seq method of Tsai et al. (2015, Nature Biotechnology 33:187–197). All model training, evaluation, and figure-generation code used in this analysis is available on request.

